# Curvature Analysis of Estrogen Receptor Positive Breast Cancer Under PI3K Inhibition

**DOI:** 10.1101/049437

**Authors:** Romeil Sandhu, Eneda Toska, Maurizio Scaltriti, José Baselga, Joseph Deasy, Jung Hun Oh, Sarah Tannenbaum, Allen Tannenbaum

## Abstract

In this note, we re-examine the work of Bosch *et al*. from a network point of view. In particular, we employ an extended defintion of Ollivier-Ricci curvature that allows us to study graphs with both positive and negative weights. This is done by utilizing a dual formulation of the Wasserstein 1-metric, allowing us to extend the Earth Mover’s Distance to signed measures. The resulting curvature may be applied study the robustness properties of general networks modelled as weighted graphs. In this note, we apply the theory to elucidate the robustness and therefore possible mechanisms of resistance of estrogen receptor positive breast cancer under PI3K inhibition.

## 1 Introduction

The present paper constitutes a re-examination of the results from [7] from a geometric network-theoretic point of view. The latter work at studied the effects of PI3Kα inhibition on estrogen receptor (ER) signaling, and at understanding its possible effect on limiting the efficacy of PI3K inhibitors. The authors found that the inhibition of the PI3K pathway triggers the activation of the ER dependent transcription machinery. This adaptive response was shown to be very important since a key consequence of the work was the observation that suppression of ER activity can sensitize tumors to PI3K inhibition.

We wish to consider these observations from a network point of view. In previous work [24], we demonstrated that a graph-theoretic notion of curvature was positively correlated to robustness defined in terms of the rate function from large deviations theory [28]. This relied on the Fluctuation theorem [9–11] in which it is demonstrated that entropy and robustness are positively correlated as well as results from [15]. More specifically in [24], we proposed an integrative framework to identify genetic features related to cancer networks and to distinguish them from the normal tissue networks by geometrical analysis of the networks provided by The Cancer Genome Atlas (TCGA) data. This relationship was exploited to show that curvature could be regarded as a *cancer hallmark*.

The underlying notion of curvature on weighted graphs is based on the Wasserstein 1-metric [19] from optimal mass transport theory [23,30,31]. This is called *Ollivier-Ricci* curvature. As such, one needs all the correlations to be positive giving well-defined positive measures in order to define the Ollivier-Ricci curvature. In the present work, based upon a dual formulation of linear programming applied to the Earth Mover’s Distance (EMD), we extend the definition of Ollivier-Ricci curvature to the more realistic case in which one allows both negative and positive weights (correlations) in our cancer networks [13,25,30,31].

We now outline the sections of the present note. In Section 2, we define the classical notion of Ricci curvature via an approach that lends itself to an extension to more general metric spaces. In Section 3, we describe the general Monge-Kantorovich problem of optimal mass transport (OMT). We state a dual formulation that allows us to treat correlation networks with both positive and negative correlations. Sections 4 and 5 are devoted to formulating the positive correlation of curvature and robustness. Then in Sections 6 and 7 Ollivier-Ricci curvature is defined on weighted graphs first in the positive weighted case, and then allowing weights to be of mixed sign. Finally, in Section 8, we analyze the results of [7] from a geometric network point of view, and then in Section 9 we outline some possible future research directions.

## 2 Background on Ricci curvature

In order to motivate generalized notions of Ricci curvature suitable for complex networks, we will begin with an elementary treatment of curvature following [12,30,31] to which we refer the interested reader for all the details. Let *M* be an *n*–dimensional Riemannian manifold, *x* ∈ *M*, let *T_x_M* denote the tangent space at *x*, and *u*_1_, *u*_2_ ∈ *T*_*x*_*M* orthonormal vectors. Then for geodesics γ_*i*_(*t*): = exp(*tu*_*i*_), *i* =1,2, the *sectional curvature K*(*u*_1_, *u*_2_) measures the deviation of geodesics relative to Euclidean geometry, i.e.,

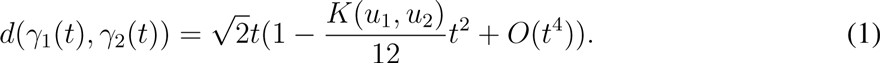

The *Ricci curvature* is the average sectional curvature. Namely, given a (unit) vector *u*∈ *T*_*x*_*M*, we complete it to an orthonormal basis, *u, u*_*2*_,…,*u*_*n*_. Then the *Ricci curvature* is defined by 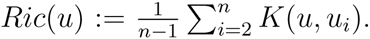. It may be extended to a quadratic form, giving the so-called *Ricci curvature tensor*.

As alluded to in the Inroduction, we will need to extend the notion of curvature to discrete graphs in order study cancer correlation networks. For general metric spaces and especially discrete spaces, ordinary notions such as differentiability necessary to define Ricci curvature as above do not make sense. There is however a very nice argument due to Villani [31] that indicates an elegant way to getting around such difficulties via the notion of convexity. Indeed, let ƒ: ℝ^n^ →ℝ.

Then if ƒ is *C*^2^, convexity may be characterized as

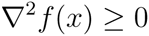

for all *x*. This is called by Villani an *analytic* definition of convexity (as the usual definition of Ricci given above). On the other hand, one can also define convexity in a *synthetic* manner via the property that

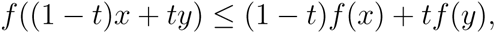

for all *x*, *y* ∈ ℝ^n^, and *t* ∈ [0,1]. In the latter case, no differentiability is necessary.

Following [15,18], one may define a synthetic notion of Ricci curvature in terms of so-called *displacement convexity* inherited from the Wasserstein geometry on probability measures. This will be detailed in Section 4 below.

## 3 Monge-Kantrovich problem

The first optimal transport (OMT) problem proposed by Monge in the 1780’s, was posed as a civil engineering problem that asks for the minimal transportation cost to move a pile of soil (“déblais”) toan excavation (“remblais”). The places where the soil should be extracted, and the ones which should be filled, are all known. A more modern form of the problem introduced by Kantorovich in 1940’s yields the so-called Monge-Kantrovich problem (MKP). The framework of the problem is as follows; see [13,30,31] for all the details. We will describe the set-up of Kantorovich.

Let (*X, μ*) and (*Y*, *ν*) be two probability spaces,and let П(*μ*,ν) denote the set of all joint probability measures on *X*×*Y* whose marginals are *μ* and *ν*. The optimal transport cost suggested by Kantorovich was the following linear programming problem:

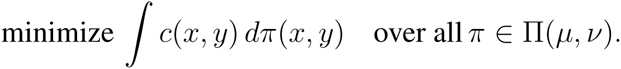

where *c*(*x, y*) is the cost for transporting one unit of mass from *x* to *y*. The cost function was originally defined in a distance form on a metric space (*X*, *d*). This leads us to the following distance function know as *L*^*p*^ Wasserstein:

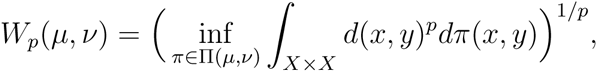

In our work, we will particularly interested in the two cases *p* = 1 and *p* = 2. The *L*^1^-Wasserstein distance is also known as the *Kantorovich-Rubinstein distance*,or *Earth Mover’s distance* among computer scientists. In [20] Ollivier used *L*^1^-Wasserstein distance to define the (Ollivier-)Ricci curvature.

We note that for *p* = 1, and for *X*, *Y* normed vector spaces, one may show that

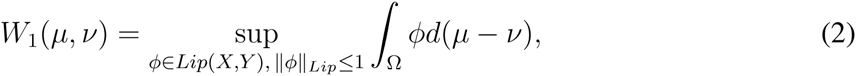

where *Lip*(*X*, *Y*)denotes the set of all Lipshitz continuous functions, and

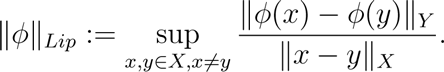

This dual formulation of the linear programming problem makes sense for signed measures, and so as we have argued in [25] may be used to formulate a notion of curvature of graphs with both positive and negative weights. This is will further explicated in Section 7 below.

## 4 Curvature and robustness

In order to elucidate the connections between the statistical mechanical notion of entropy, and the geometric notion of curvature, we sketch some of the work of Lott and Villani [15].

Let (*X, d, m*) denote a geodesic space, and set

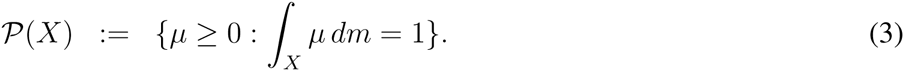

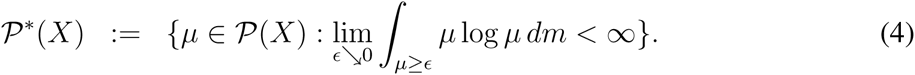

We define

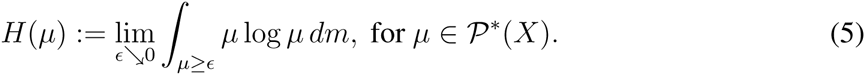

The latter is the negative of the *Boltzmann entropy S*(*μ*):=— H(*μ*); note that the concavity of *S* is equivalent to the convexity of *H*. In the classical case when *X* is a Riemannian manifold, one may prove that *X* has Ricci curvature bounded from below by *k* if and only if for every *μ*_0_, *μ*_1_ ∈ *p*^*^ (*X*), there exists a constant speed geodesic *μ*_*t*_ with respect to the Wasserstein metric connecting *μ*_*0*_ and *μ*_1_ such that

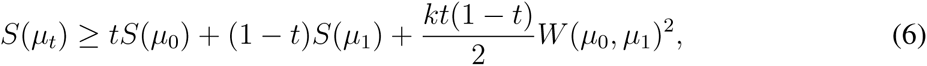

for 0 ≤ *t* ≤ 1. This means that changes in entropy and curvature are ***positively correlated***. We express this relationship as

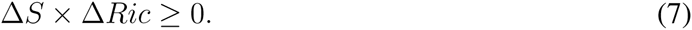

In the general geodesic space case, (6) is taken to be the ***definition*** of Ricci curvature.

We will describe a specific notion of Ricci curvature and entropy on weighted graphs below. We just note here that changes in *robustness*, i.e., the ability of a system to functionally adapt to changes in the environment (denoted as ΔR) is also positively correlated with entropy via the Fluctuation Theorem [9–11], and thus with network curvature:

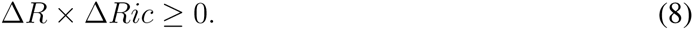

See the next section for a discussion of the Fluctuation Theorem. Since the curvature is very easy to compute for a network as we will see in Section 6, this may be used as an alternative way of expressing functional robustness.

## 5 Fluctuation theorem

We give now an informal description of the Fluctuation Theorem [9–11]. Letting *p*_∈_(*t*) denote the probability that the mean of a given network observable deviates by more thane from the original (unperturbed) value at time *t*,we set

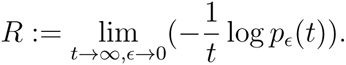

This is the *rate function* from large deviations theory [28].

On the other hand, evolutionary entropy *S* may be characterized in this setting as

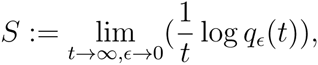

where *q*_∈_(*t*) denotes the minimal number of genealogies of length *t* whose total probability exceeds 1 — ∈. Thus the greater the *q*_∈_(*t*),the smaller the *p*_∈_(*t*) and so the larger the decay rate. The Fluctuation Theorem [9–11] is an expression of this fact for networks, and can be expressed as

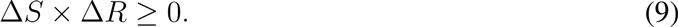

Considering (6), we conclude that changes in robustness *(ΔR)* is also positively correlated with the network curvature, as stated in (8). This latter relationship will be a key in studying the robustness of cancer networks.

## 6 Ollivier-Ricci curvature: positive weights

In this section, we will sketch an elegant approach to Ricci curvature due to Ollivier [19, 20] that seems ideal for studying network robustness in many practical situations given its ease of computation. The approach was developed as a discrete analogue of a defining property of Ricci curvature in the continuous case. The idea is that for two very close points *x* and *y* on a Riemannian manifold with respective tangent vectors *w* and *w’*, in which *w’* is obtained by a parallel transport of *w*, the two geodesics will get closer if the curvature is positive. This is reflected in the fact that the distance between two small (geodesic balls) is less than the distance of their centers. Ricci curvature along direction *xy* reflects this, averaged on all directions *w* at *x*. Similar considerations apply to negative and zero curvature [32].

Now let (*X, d*) denote a geodesic metric space equipped with a family of probability measures {*μ*_*x*_: *x* ∈ *X*}. Following [19,20], we define the *Olliver–Ricci curvature K*(*x*, *y*) along the geodesic connecting *x* and *y* via

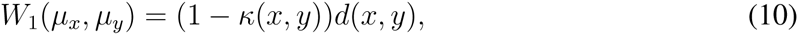

where *W*_1_ denotes the Earth Mover’s Distance (Wasserstein 1–metric), and *d* the geodesic distance on the space. For the case of weighted (undirected) graphs of greatest interest in networks, we set

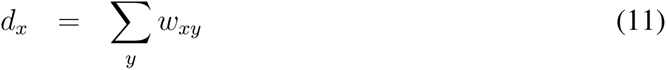

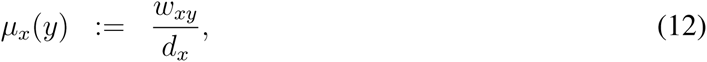

the sum taken over all neighbors of *x* where *w*_*xy*_ denotes the weight of an edge connecting *x* and *y* (it is taken as zero if there is no connecting edge between *x* and *y*). ***All the interaction weights here are positive***. The measure *μ*_*x*_ may be regarded as the distribution of a one-step random walk starting from *x*. As is argued in [19], this definition is more inspired from an approach such as that given via equation (1). An advantage of this, is that it is readily computable since the Earth Mover’s Metric may be computed via linear programming [13,23,31].

## 7 Ollivier-Ricci curvature: positive and negative weights

The correlation networks that we will be considering from cancer have both positive and negative weights, and so one needs a notion of curvature for this case as well, i.e. for weighted undirected graphs with weights *w*_*xy*_ that may be either positive and negative. Accordingly, we need an extension of the Wasserstein distance for signed measures. See also [16] for an extensive discussion about various possible approaches for Wasserstein on signed measures. Approaches to the extension of Ollivier–Ricci curvature to very general networks modelled by graphs with both positive and negative weights in both the directed and undirected cases may be found in [25].

In our case, we note that as in Section 3, using duality, one can show that

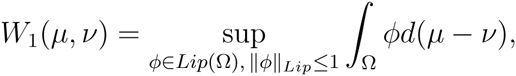

and since this characterization of the Wasserstein 1–metric only depends on the difference of the two measures, it is valid for signed measures, in particular graphs with both positive and negative weights.

In the discrete case, using the above notation, for weighted graphs we have that

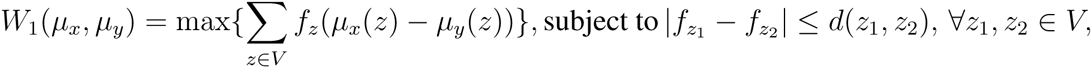

which is clearly a linear programming problem [13]. Again this makes sense for graphs with both positive and negative weights. Here *μ*_*x*_ and *μ*_*y*_ are signed probability measures defined exactly as in Equation 11 above.

Finally, following [19], we define the Olliver–Ricci curvature *K*(*x, y*) via:

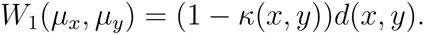

We now apply this to the analysis of an important cancer network.

## 8 Results: ER+ breast cancer under PI3K inhibition

One of the main attractions with the utilization of weighted graphs whose interactions (edges) are given by signed measures, lies in their application to cancer correlation networks, and the potentialof better elucidating mechanisms of drug targeting [24, 33]. In this section, we report our preliminary results concerning the application of our discrete geometric approach to the previous work of [7] in order to try to identify mechanisms of resistance. We should note that the networksexamined do not take into account any notion of directionality (e.g., inhibitory and activator effects) and the underlying topology represented a protein–to–protein interaction network. This was constructed using *clinically proven* links derived from the Human Protein Reference Database (HPRD).

We should emphasize that the work presented here is preliminary, but does provide further supporting evidence for the conclusions of [7]. Indeed, this network approach does seem to give a different point of view to better the understanding of drug resistance. To this end, Table 1 provides global curvature results as well as the expected value of the left tail (ELT) of the distribution (similar to the notion of p–values) to highlight an approximate lower bound of Ricci curvature. We see that at the global network level, the treatment of BYL710 inhibitor for PI3K first induces fragility prior to an increase in robustness before such resistance subsides over a 48-hour time-window. Moreover, we examined specific genes seen in Figure 1. As noted in [7], during initial treatment, the activity of the genes exhibits fragility prior to building resistance and then subsequently subsides. This is in line with gene expression data where maximal expression of ER–related genes is seen at the 24 hour mark and then subsequently subsides. Interestingly, this effect seems to be greatest on PI3KR1, which makes biological sense since we are considering the effect of a PI3K inhibitor. In this case, we see a very large increase in fragility at 4 hours (exhibited by large negative curvature). All together, such results begin to point towards the notion of utilized network geometry and in general, network modeling, to allow for one to assess biological drug sensitivity in a more quantitative non–subjective manner.

**Table 1:**
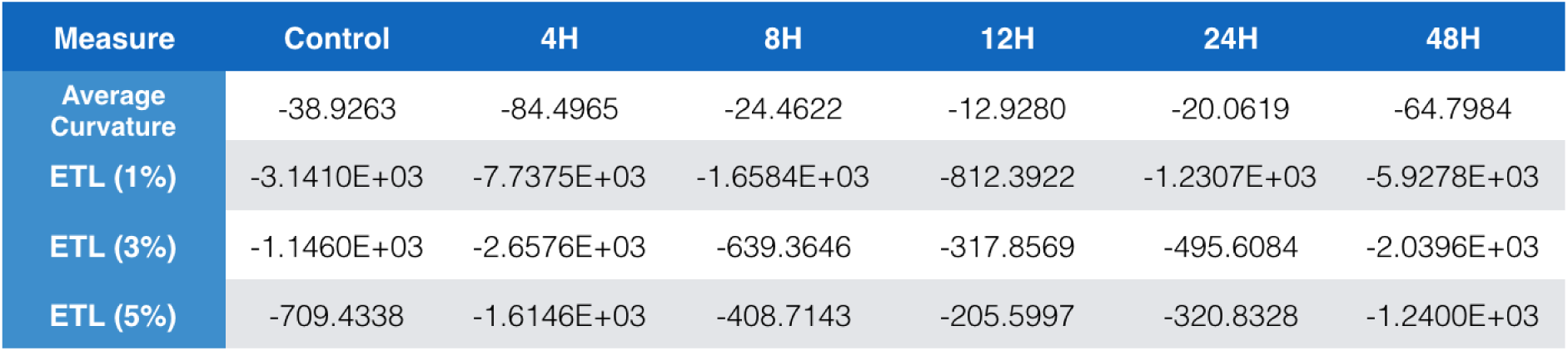
This table presents average curvature results (in the sense of Ricci curvature) across the time-varying data first presented in [7].

| Measure | Control | 4H | 8H | 12H | 24H | 48H |
| --- | --- | --- | --- | --- | --- | --- |
| Average Curvature | -38.9263 | -84.4965 | -24.4622 | -12.9280 | -20.0619 | -64.7984 |
| ETL (1%) | -3.1410E+03 | -7.7375E+03 | -1.6584E+03 | -812.3922 | -1.2307E+03 | -5.9278E+03 |
| ETL (3%) | -1.1460E+03 | -2.6576E+03 | -639.3646 | -317.8569 | -495.6084 | -2.0396E+03 |
| ETL (5%) | -709.4338 | -1.6146E+03 | -408.7143 | -205.5997 | -320.8328 | -1.2400E+03 |

**Figure 1:**
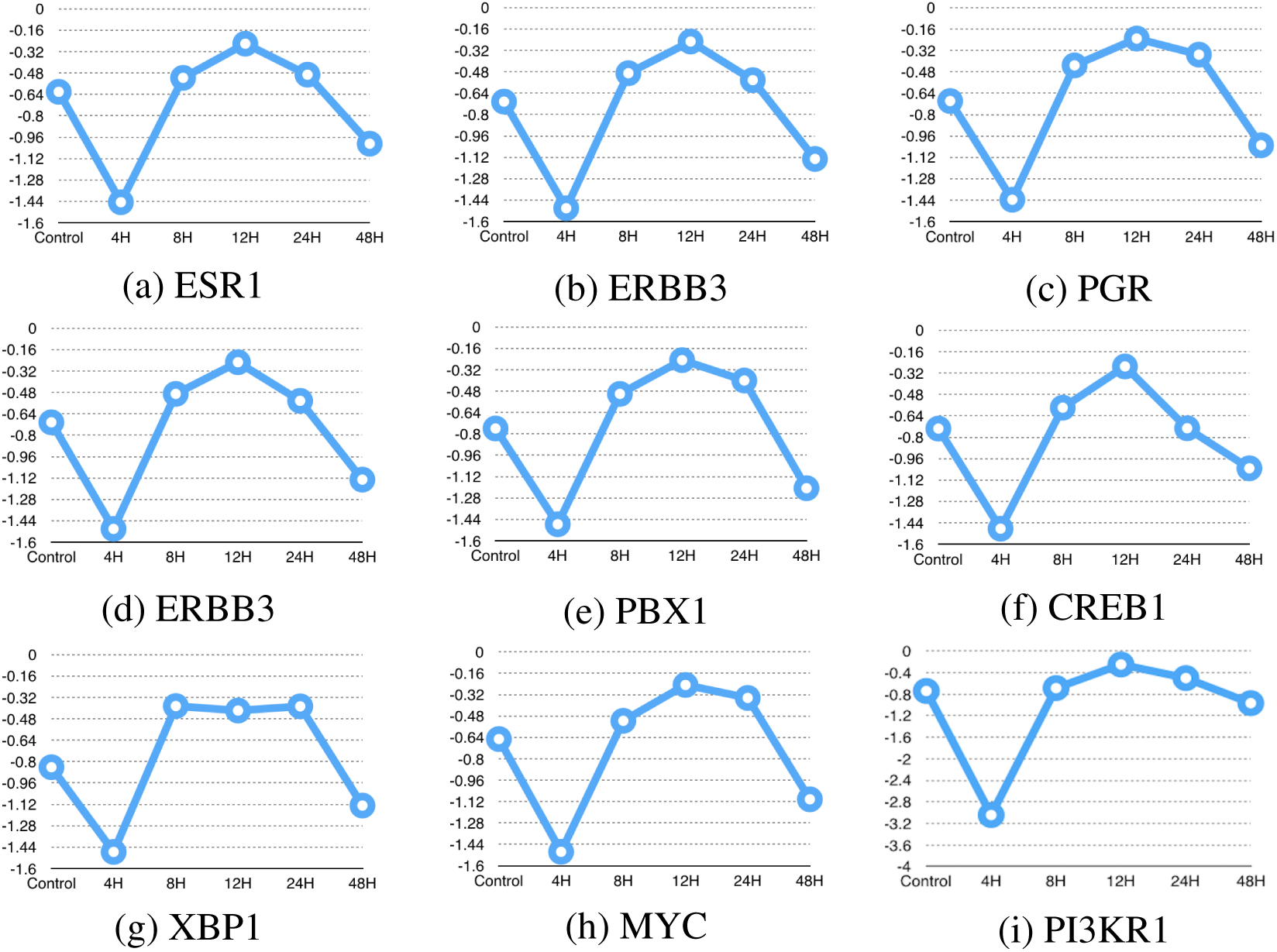
Summation of Ricci curvature of all direct and indirect pathways. As noted above, during initial treatment, ER–related genes and activity exhibits fragility prior to building resistance and then subsequently subsides. Note: Results are presented minus scaling factor of 1e3 for figures (a)–(h) and 1e5 for figure (i).

## 9 Future Work & Summary

In this paper, we employed an extension of Olliver-Ricci curvature [19,25] valid for rather general networks in order to further our understanding of the robustness and fragility of biological cancer (correlation) networks. This was explicitly illustrated via the preliminary results on the robustness of certain breast cancer networks following the work of [7].

Future work will be built around using these geometric ideas to try to build models of resistance via the identification of key hubs of resistance (robustness as indicated by high curvature) and drugsensitivity (fragility as indicated by lower curvature). Inhibition and activation will also need to be modeled and analyzed through the directed methodology [25].

